# SARS-CoV-2 ORF8 modulates the upper respiratory tract inflammatory response to facilitate transmission

**DOI:** 10.64898/2026.08.09.743817

**Authors:** Grace O. Ciabattoni, Stacey Bartlett, Marisa E. McGrath, Matthew B. Frieman, Meike Dittmann, Mila B. Ortigoza

## Abstract

The ability of respiratory viruses to exploit host immune responses to promote transmission is a defining feature of pandemics. SARS-CoV-2 remains a major global public health threat because of its persistent evolution and capacity to counteract evolving immune defenses. Although the immune evasion properties of the SARS-CoV-2 Spike protein are well characterized, the contributions of other viral proteins to transmission remain poorly understood. Here, we used the infant mouse model to define the role of the accessory protein ORF8 in SARS-CoV-2 spread. We demonstrate that ORF8 supports efficient upper respiratory tract (URT) infection, infectious virus shedding, and host-to-host transmission. Mice infected with a recombinant SARS-CoV-2 strain lacking ORF8 (rΔORF8) had less infectious virus recovered from URT tissues and nasal secretions and transmitted less efficiently than mice infected with the isogenic ancestral strain rWA-1, which contains an intact ORF8. Recombinant viruses encoding naturally occurring ORF8 mutations exhibited distinct transmission phenotypes, with ORF8-deficient viruses resembling rΔORF8. Infection with ORF8-sufficient viruses induced greater macrophage recruitment, inflammatory cytokine production, and type-I interferon (IFN-I) signaling programs than ORF8-deficient viruses. Intranasal IFNβ supplementation partially restored URT shedding by rΔORF8-infected mice and rescued transmission to contacts, whereas blockade of the type I interferon receptor (IFNAR) in rWA-1-infected index mice reduced contact infection and transmission. Together, these findings demonstrate that ORF8 promotes SARS-CoV-2 transmission by engaging an IFN-I-associated inflammatory and secretory program in the URT that supports virus shedding from the infected host. These data identify ORF8 as a viral determinant of host mucosal responses that promote contagiousness.

**Importance:** Efficient host-to-host transmission underlies the success of respiratory viruses. Although SARS-CoV-2 research has largely focused on the Spike protein, accessory proteins can also shape viral fitness and spread. We previously identified ORF8 as a determinant of SARS-CoV-2 transmission. Here we show that ORF8 promotes SARS-CoV-2 infectious viral shedding and transmission by engaging IFN-I-associated inflammatory and secretory responses in the URT. These findings reveal how a SARS-CoV-2 accessory protein can exploit mucosal antiviral responses to increase host contagiousness.

## Introduction

Since its emergence in late 2019, SARS-CoV-2 has remained a persistent and evolving threat to global public health. Its rapid evolution has sustained global spread and enabled immune escape from several antibody-based interventions, including monoclonal antibodies and vaccine-induced neutralizing responses (1–4). Developing effective countermeasures against SARS-CoV-2 requires deeper understanding of viral factors beyond the Spike protein and host responses that influence transmission.

Transmission of respiratory viruses is a multistep process. The virus must establish infection in the upper respiratory tract (URT) of an infected host (index), be released (shed) into respiratory secretions, and subsequently establish infection in a susceptible host (contact) (5).

Transmission is favored when the timing, proximity, and duration of exposure are sufficient to deliver an infectious dose to a susceptible host. These variables are influenced by the inflammatory and secretory state of the URT, which determines both the amount of infectious virus available for expulsion and the efficiency with which it is released into the environment. The URT is the primary site of SARS-CoV-2 infection and has a dual role in respiratory defense and transmission (6). It functions as a frontline barrier that limits pathogen spread to the lower respiratory tract (LRT), but it also serves as the principal site from which virus is released into the environment (6). Mucosal defenses trap inhaled pathogens in respiratory secretions and facilitate their clearance through nasal discharge, mucociliary transport, coughing, sneezing, and talking. These responses are antiviral because they help clear virus from the infected host. For respiratory viruses, however, the same processes can increase the release of infectious virus and thereby enhance contagiousness.

The URT differs from the LRT in its microbiota, local conditions, epithelial composition, and immune architecture (7), producing compartment-specific immune responses and divergent clinical outcomes. URT inflammation activates innate antiviral pathways, including type-I interferon (IFN-I) signaling, and is often accompanied by symptoms such as rhinorrhea and congestion. By increasing mucus production and triggering coughing and sneezing, these responses trap virus in respiratory secretions and promote its expulsion into the environment (8–10). In contrast, LRT inflammation is more closely associated with disease severity, including pneumonia, acute respiratory distress syndrome (ARDS), excessive cytokine release (cytokine storm), and impaired gas exchange (11). Therefore, immune activation in the URT and LRT can have distinct consequences for viral fitness: URT inflammation shapes contagiousness, whereas LRT inflammation drives pathogenesis.

The timing and magnitude of URT infection and host responses are critical to transmission efficiency. Rapid accumulation of infectious virus in nasal secretions correlates with increased transmission (12). Efficient spread therefore requires viral strategies that preserve infection in the URT while selectively engaging host responses that promote the release of virus-containing secretions. SARS-CoV-2 accessory proteins are well positioned to influence this balance because they remodel host pathways without serving as the primary receptor-binding protein.

These include ORF3a, which can modulate autophagy and inflammatory signaling (13), ORF6, which antagonizes IFN-I signaling (14), and ORF7a, which has been linked to antagonism of the restriction factor BST-2/Tetherin (15, 16). Among these accessory proteins, ORF8 is notable for its rapid evolution and diverse immune-modulatory functions.

ORF8 has evolved extensively across SARS-CoV-2 lineages and shares limited sequence homology with its counterpart in SARS-CoV-1 (17–19). The SARS-CoV-2 ORF8 gene encodes a 121 amino acid (AA) protein with an N-terminal signal sequence that directs endoplasmic reticulum (ER) import and a C-terminal immunoglobulin (Ig)-like fold (20–22). ORF8 can also be secreted and detected in the serum of infected patients (20, 23). It has been reported to downregulate major histocompatibility complex class I (MHC-I) to reduce cytotoxic T cell lymphocytes (CTLs) recognition (24); bind to the Fc receptor CD16a on monocytes and NK cells to impair antibody-dependent cellular cytotoxicity (ADCC) (25); and act as a cytokine-like protein (virokine) that perturbs immune signaling and induces inflammatory cytokine production (26, 27). Consistent with these functions, human infection with an ORF8-deletion variant was associated with dampened inflammation, including diminished IFN-associated responses, and milder COVID-19 disease (28). Together, these observations suggest that ORF8 can reshape host defenses in ways that may influence viral persistence, mucosal inflammation, and transmission.

Despite extensive characterization of ORF8’s immune-modulatory functions, its role in transmission remains incompletely defined. Building on our previous study that identified ORF8 as a key determinant of SARS-CoV-2 transmission in the infant mouse model (29), we focused on the URT as the compartment where viral fitness, mucosal inflammation, secretion biology, and host-to-host spread converge. Mice infected with an ORF8-deleted SARS-CoV-2 strain exhibited lower infectious virus in nasal secretions and reduced transmission compared to its isogenic, ORF8-sufficient wild-type control, suggesting that ORF8 supports contagiousness by shaping the URT inflammatory environment that promotes viral expulsion. Here, we define how ORF8 influences URT inflammation, immune cell recruitment, IFN-I-associated signaling, infectious virus shedding, and host-to-host transmission.

## Results

### Naturally occurring ORF8 variants reveal distinct transmission phenotypes

Our previous work identified ORF8 as a factor that enhances SARS-CoV-2 transmission in the infant mouse model by increasing infectious virus recovered from URT tissues and secretions (29). ORF8 is one of the most variable genetic loci in SARS-CoV-2, with lineage-associated mutations observed in several variants of concern (VOCs) and circulating viruses (**Table 1**). The ancestral SARS-CoV-2 strain first isolated in Wuhan, China (Wuhan-1), was classified into “L” or “S” strains based on amino acid (AA) 84 in ORF8 (30). The S84L mutation has since been observed in multiple SARS-CoV-2 lineages. The Gamma variant carried both S84L and E92K mutations in ORF8, suggesting potential functional adaptation. ORF8 truncations and deletions have also emerged. The Alpha variant carried a premature stop codon at AA 27 producing an ORF8 protein truncated by approximately 75% (Q27*) (21), while later Omicron lineages have recurrently lost ORF8 expression through mutations such as G8* in XBB1.5 or larger deletions spanning ORF7a, ORF7b, and ORF8 in BA.3.2 (2, 3). Alpha and Omicron were highly fit viruses in humans however, their transmission success occurred in the context of extensive variant-defining mutations in the Spike protein that enhanced receptor binding, immune evasion, or both, potentially masking the contribution of accessory proteins to viral fitness (31–33). As a result, the specific contribution of ORF8 to transmission has remained difficult to resolve from epidemiologic data alone.

**Table 1:**
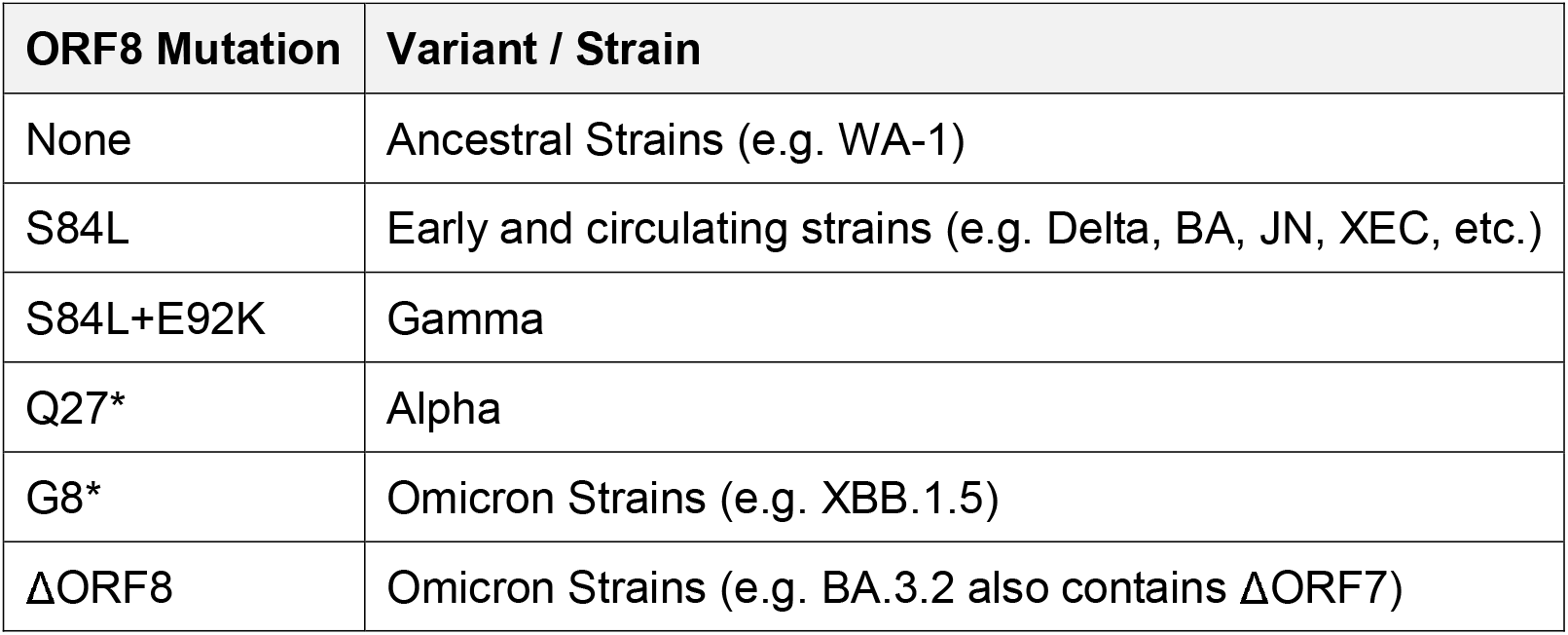
Natural ORF8 Mutations and their Associated SARS-CoV-2 Variants.

Naturally occurring ORF8 mutations provide important biological context for understanding the role of ORF8 in SARS-CoV-2 transmission. To directly evaluate the isolated effects of these mutations on transmission efficiency, we generated a panel of recombinant SARS-CoV-2 viruses encoding representative ORF8 variants in the wild-type SARS-CoV-2 strain isolated in Washington state (rWA-1, SARS-CoV-2 USA_WA-1/2020). This panel included viruses lacking ORF8 (rΔORF8) or ORF7a (rΔORF7a), as well as viruses encoding ORF8 point mutations: S84L, S84L+E92K, E92K, and Q27* (**Fig.1A**) (32, 33). Using the established infant mouse model of transmission with K18-hACE2^(+/-)^ mice, we targeted URT infection by intranasal instillation of 1500 plaque-forming units (PFU) of rSARS-CoV-2 in a low-volume inoculum (5, 29, 34). Following infection, index mice were co-housed with their dam and naïve littermates (contacts), and cohorts were monitored for 7 days. Viral shedding from both index and contact mice was collected daily by gently dipping each pup’s nares into viral medium, and expelled infectious virus was quantified by plaque assay (**Fig.1B**). This shedding assay provides a longitudinal measure of infectious virus recovered from luminal URT secretions and serves as a proxy for transmission potential.

**Figure 1.**
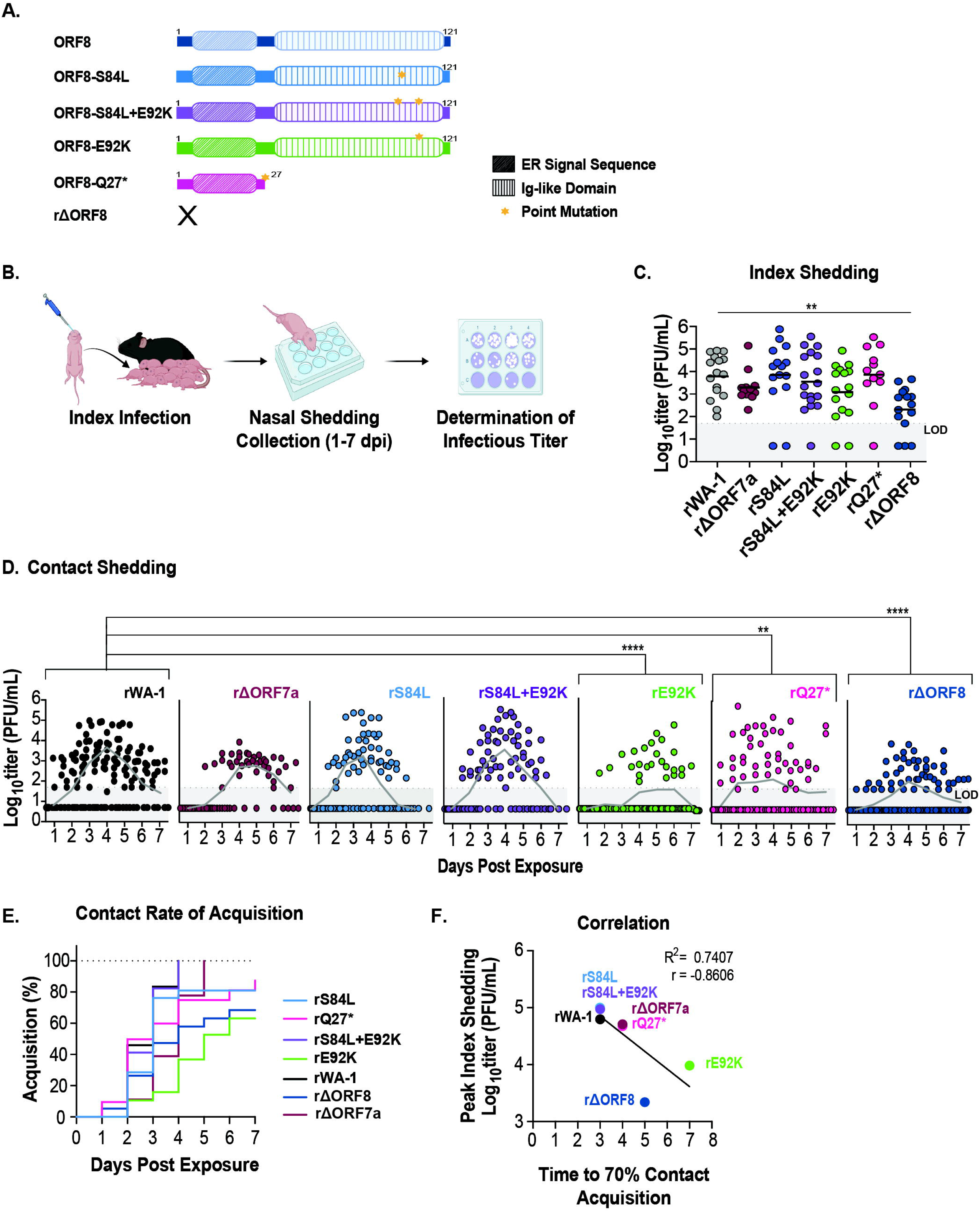
SARS-CoV-2 ORF8 variants shape transmission dynamics. (A) Cartoon of ORF8 mutations in the recombinant virus panel. Mutation sites are indicated by yellow stars, and domain architecture is shown. (B) Schematic of the infant mouse transmission model. Index pups were intranasally inoculated on day 0 and co-housed with naïve contact pups for 7 days. Nasal shedding was collected daily from index and contact pups separately and titrated by plaque assay. (C) Index shedding titers collected on days 1–3 post infection were pooled within each viral group and compared across the virus panel. (D) Contact shedding titers collected on days 1-7 are plotted by day post exposure to show the temporal distribution of viral acquisition, whereas statistical comparisons pooled all contact shedding samples within each virus group across the full sampling period rather than analyzing individual days separately. (E) Transmission is plotted as percent acquisition, with onset scored as the first day of detectable infectious virus shedding. (F) Correlation between peak index shedding titers and time to 70% acquisition in contacts. <u>Statistics:</u> (C-D) Each point represents an individual shedding sample. Geometric means are shown by the horizontal black line in (C) and by gray lines in (D). Differences among viral groups in (C) and (D) were analyzed using a Kruskal–Wallis test followed by Dunn’s multiple-comparisons test, with rWA-1 as the reference group. Values below the limit of detection (LOD; 50 PFU/mL) were included and assigned a value of 5 PFU/mL, one log below the LOD. Experiments represent ≥2 independent biological replicates. Significance is denoted as *p < 0.05, **p < 0.01, ***p < 0.001, and ****p < 0.0001. Cartoons were created using BioRender.com. Graphs generated using GraphPad Prism 10.

Index shedding titers pooled across 1–3 days post-infection (dpi) were not statistically different to rWA-1 across the panel, except for rΔORF8, which shed at significantly lower levels (**Fig.1C**). Because rΔORF8 differs from rWA-1 only by the absence of ORF8, these data indicate that ORF8 is specifically required to sustain high levels of infectious virus in URT secretions.

Contact shedding further supported a specific role for ORF8 in transmission. Contacts exposed to ORF8-sufficient viruses, including rWA-1, rΔORF7a, rS84L, and rS84L+E92K, generally shed at higher titers than contacts exposed to ORF8-deficient viruses, including rΔORF8 and rQ27* (**Fig.1D**). However, individual ORF8 mutations produced distinct phenotypes. rS84L+E92K supported efficient contact shedding and acquisition, whereas rE92K was associated with reduced contact shedding despite encoding an intact ORF8 protein. This suggests that individual ORF8 mutations can alter transmission-relevant phenotypes in ways that are not explained solely by ORF8 presence or absence.

Cumulative acquisition further stratified the virus panel (**Fig.1E**). Contacts exposed to ORF8-sufficient viruses (rWA-1, rΔORF7a, and rS84L+E92K) reached 100% acquisition, whereas contacts exposed to ORF8-deficient viruses (rΔORF8 and rQ27*) reached incomplete final acquisition of 77% and 88%, respectively. Notably, two ORF8-sufficient point mutants also showed incomplete acquisition: rS84L reached 80%, and rE92K reached 60%. These data indicate that loss or truncation of ORF8 impairs transmission-relevant phenotypes, while individual ORF8 point mutations can produce distinct intermediate phenotypes rather than simply behaving as ORF8-sufficient or ORF8-deficient viruses. As in our prior studies (29, 35), peak index shedding titers at 2 dpi correlated with transmission efficiency across the panel (**Fig.1F**). These findings show that ORF8 variation shapes infectious virus shedding and contact acquisition, with ORF8 loss or truncation reducing transmission-relevant phenotypes and individual ORF8 point mutations producing distinct effects.

### SARS-CoV-2 ORF8 selectively promotes infectious virus recovered from the URT

To determine whether the reduced transmission observed in ORF8-deficient viruses reflected differences in infectious virus across respiratory compartments, we quantified viral titers at 2 dpi, the time of peak URT shedding. We collected shedding samples, representing infectious virus recovered from nasal secretions; nasal lavages, representing infectious virus recovered from URT tissue; and lung samples, representing infectious virus recovered from the LRT tissue (**Fig.2A**). In rWA-1-infected mice, infectious titers in both nasal secretions and URT tissues were significantly higher than in rΔORF8-infected mice, while lung titers were not significantly different between groups (**Fig.2B**). This pattern argues against a uniform reduction in infectious virus across respiratory compartments. Instead, ORF8 deficiency produced a selective reduction in infectious virus recovered from URT tissues and nasal secretions.

**Figure 2.**
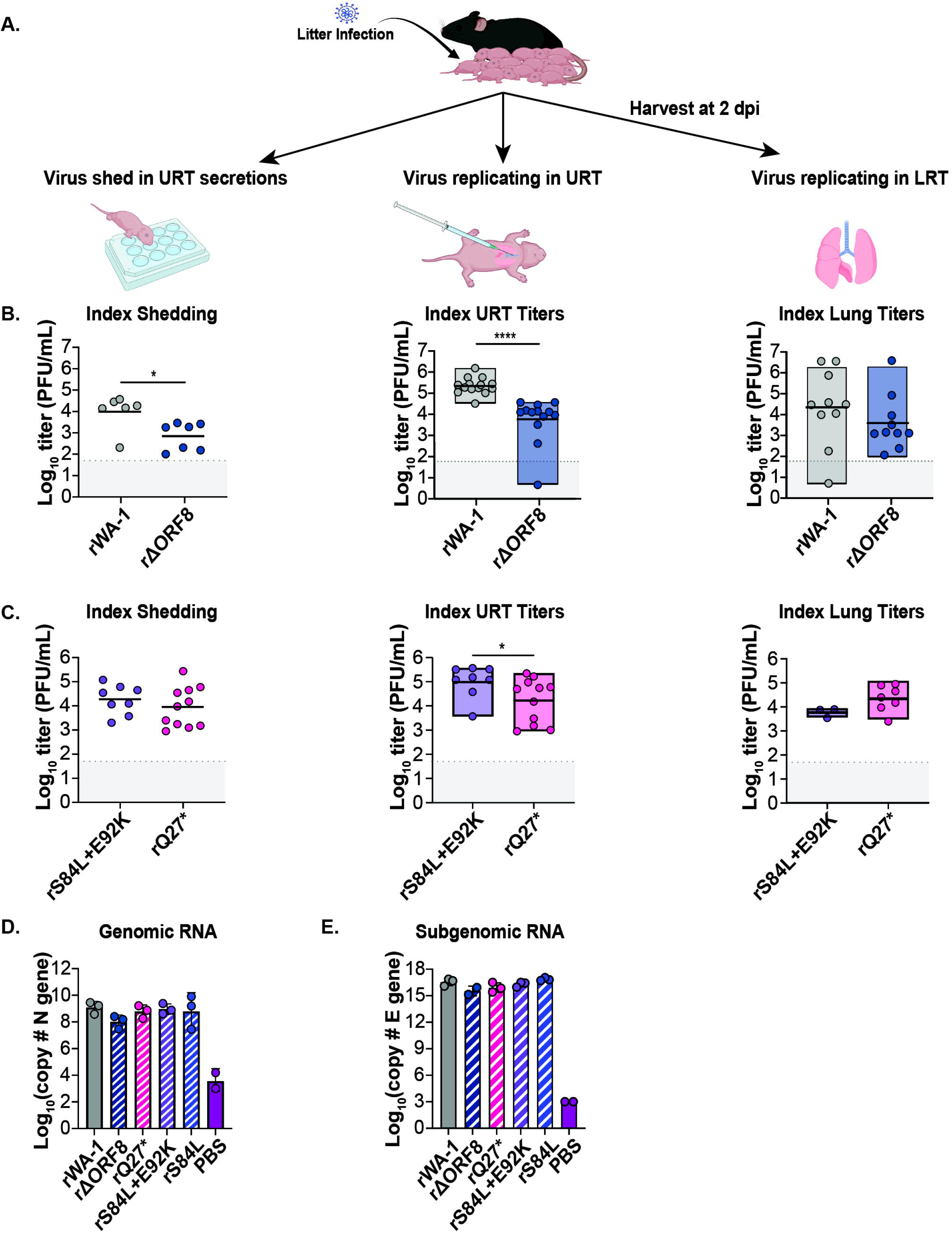
SARS-CoV-2 ORF8 promotes URT infection and shedding. (A) Experimental schematic. Index pups were inoculated with rWA-1, rΔORF8, rS84L+E92K, rQ27*, or PBS. Nasal shedding was collected at 2 dpi, and URT tissue lavages and lungs were harvested immediately afterward. (B-C) Infectious virus titers in nasal shedding samples, URT tissues (from nasal lavages after euthanasia), and LRT tissues (lungs) are shown from left to right. (B) rWA-1 versus rΔORF8. (C) rS84L+E92K versus rQ27*. (D-E) RT-qPCR quantification of viral RNA in URT tissues: (D) N gene genomic RNA and (E) E gene leader-TRS subgenomic RNA. Viral RNA copy number was determined using experiment-specific standard curves. <u>Statistics:</u> Each point represents an individual mouse, and geometric means are shown. Differences between two groups in (B) and (C) were analyzed using a Mann-Whitney U test. Figures 2D and 2E were analyzed using a Kruskal-Wallis test followed by Dunn’s multiple-comparisons test, with rWA-1 as the reference group. No statistical significance was identified. Values below the LOD for infectious virus titers were assigned a value of 5 PFU/mL, one log below the LOD. Experiments represent ≥2 independent biological replicates. Significance is denoted as *p < 0.05, **p < 0.01, ***p < 0.001, and ****p < 0.0001. Cartoons were created using BioRender.com. Graphs generated using GraphPad Prism 10.

We next considered whether this URT phenotype could reflect the pattern of ACE2 expression in our mouse model strain, K18-hACE2^(+/-)^, in which human ACE2 (hACE2) is driven by the keratin 18 promoter and broadly distributed across epithelial cells (36–38). To address this, we repeated the analysis in hACE2-knock-in (KI) mice, in which hACE2 replaces the mouse Ace2 at the endogenous locus and is expressed under physiological regulatory control (31). The URT-restricted reduction in rΔORF8 titers was recapitulated in this second mouse model (**Supplemental Fig.1A)**, indicating that the rΔORF8 phenotype is not explained by the transgenic ACE2 expression pattern of K18-hACE2 mice.

We then examined representative recombinant ORF8-mutant viruses for their effects on URT and LRT infectious titers. The ORF8-deficient truncation mutant rQ27* yielded significantly lower URT titers, resembling rΔORF8, whereas the ORF8-sufficient virus rS84L+E92K produced higher URT titers, resembling rWA-1. Lung titers remained comparable across viruses (**Fig.2C**). These data extend the compartment-specific rΔORF8 phenotype to an independently generated ORF8 truncation mutant.

To assess whether differences in infectious virus recovered from nasal secretions and URT tissues were accompanied by differences in viral RNA production, we quantified viral transcripts at 2 dpi in URT compartments using TaqMan PCR. Genomic RNA (gRNA) was measured by nucleocapsid (N) gene copy number. Although gRNA levels trended slightly lower in rΔORF8 samples, no significant differences were detected between viruses (**Fig.2D**). To examine active viral transcription, we quantified subgenomic RNA (sgRNA) for the envelope (E) gene containing the leader transcriptional regulatory sequence (TRS). No statistically significant differences in sgRNA levels were observed across viruses (**Fig.2E**). These findings indicate that ORF8 promotes recovery of infectious virus from URT tissues and nasal secretions without a statistically significant change in genomic or subgenomic RNA levels. This distinction provided the rationale for testing whether ORF8 shapes URT host responses associated with infectious virus shedding.

### ORF8 augments inflammatory and type-I-interferon-associated responses in the URT

Respiratory viral infections trigger localized inflammatory responses in the URT that serve multiple host-defense functions, including restricting spread to the LRT and promoting viral clearance through mucus production, mucociliary transport, and epithelial defense programs (10, 39–41). These responses can reduce viral burden within the infected host while simultaneously increasing the amount of infectious virus released into the environment. Based on the compartment-specific reduction in infectious virus observed with rΔORF8, we hypothesized that ORF8-sufficient viruses would induce a stronger inflammatory and secretory URT state associated with enhanced infectious virus shedding.

To test this, infant mice were inoculated with PBS or recombinant SARS-CoV-2 viruses, and nasal lavages were collected at 2 dpi to quantify cytokine and chemokine responses by Luminex. Compared with rΔORF8-infected mice, rWA-1-infected mice exhibited significantly higher levels of inflammatory mediators associated with antiviral innate immune activation, including MIP-1α, MIP-1β, RANTES, and IL1α (**Fig.3A, Supplemental Fig. 2A)**. In contrast, rΔORF8 infection elicited substantially lower levels of these inflammatory mediators. A similar ORF8-dependent pattern was observed when comparing naturally occurring ORF8 variants: the ORF8-sufficient virus rS84L+E92K induced higher cytokine and chemokine levels than the ORF8-deficient truncated mutant rQ27* (**Fig.3A)**. Together, these findings associate ORF8 sufficiency with stronger URT inflammation and greater recovery of infectious virus from URT tissues and nasal secretions.

**Figure 3.**
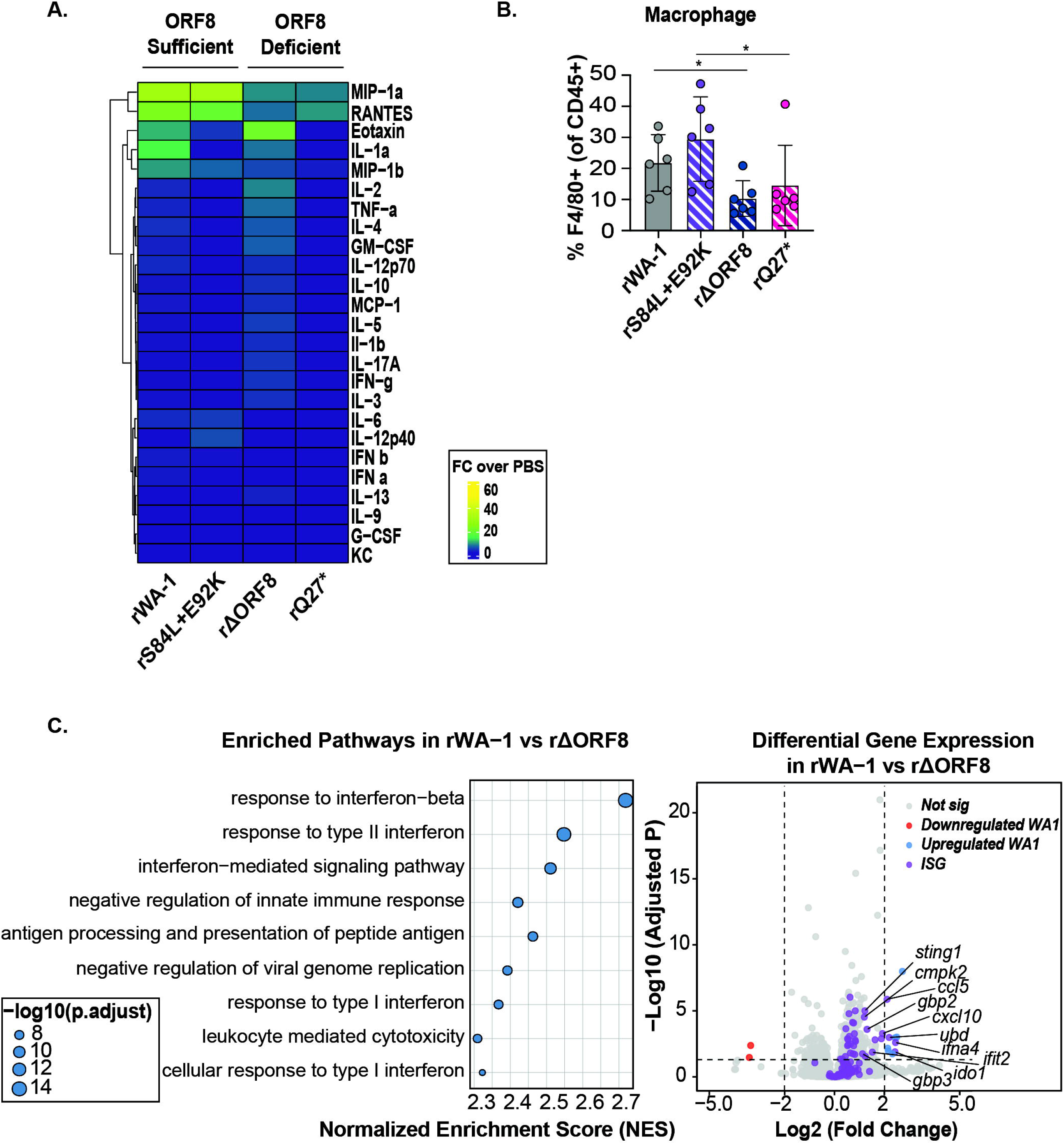
SARS-CoV-2 ORF8-sufficient viruses enhance URT inflammation. Infant mice were infected with the indicated recombinant SARS-CoV-2 viruses, euthanized at 2 dpi, and nasal lavages were collected for inflammatory profiling. (A) Heatmap showing cytokine and chemokine levels measured by Luminex in nasal lavages from mice infected with rWA-1, rS84L+E92K, rΔORF8, or rQ27*. Values were normalized to PBS-treated controls, and clustering was performed in RStudio. (B) Flow cytometry analysis of macrophage abundance in nasal lavage samples from rWA-1 versus rΔORF8 and rS84L+E92K versus rQ27* infection. Samples were acquired on a Bio-Rad ZE5 Cell Analyzer and analyzed in FlowJo. (C) Bulk RNA-seq analysis of nasal lavage cellular fractions from infant pups infected with rWA-1 or rΔORF8 and harvested at 2 dpi. Differentially expressed genes (DEG) were identified, and Gene Set Enrichment Analysis (GSEA) was performed. Left: GSEA of rWA-1 versus rΔORF8. Right: DGE of rWA-1 versus rΔORF8. Significant genes are indicated in blue for genes upregulated in rWA-1 and red for genes downregulated in rWA-1. Interferon-stimulated genes are indicated in purple. <u>Statistics:</u> Each point represents an individual mouse. Flow cytometry data in Figure 3B were analyzed using a Mann-Whitney U test. Experiments represent ≥2 independent biological replicates. Significance is denoted as *p < 0.05, **p < 0.01, ***p < 0.001, and ****p < 0.0001. Graphs were generated using RStudio and GraphPad Prism 10.

Cytokine and chemokine signaling regulates immune cell recruitment to sites of infection. To determine whether ORF8 influences the immune cell composition of the URT, we analyzed nasal lavage cells from infected mice by flow cytometry at 2 dpi. rWA-1 infection altered leukocyte recruitment compared to rΔORF8 infection, including a modest increase in total myeloid cells (**Supplemental Fig.2B**). Further characterization of myeloid subsets revealed enrichment of macrophages in rWA-1-infected mice (**Fig.3B**), while neutrophil and eosinophil levels were not significantly different between groups (**Supplemental Fig.2C-D**). A similar pattern was observed when comparing rS84L+E92K and rQ27* infections, with rS84L+E92K showing increased macrophage recruitment relative to rQ27* infection (**Fig.3B**).

To determine whether ORF8-associated URT inflammation was accompanied by transcriptional remodeling, we performed bulk RNA sequencing on nasal lavage samples collected from mice infected with either rWA-1 or rΔORF8. Differential gene expression was analyzed by Gene Set Enrichment Analysis (GSEA). IFN-associated pathways were among those most strongly enriched during rWA-1 infection compared with rΔORF8 infection, including responses to IFN-I, particularly IFN beta (IFNβ), and IFN-mediated signaling **(Fig. 3C)**. This transcriptional pattern coincided with higher IFNβ levels in nasal lavage fluid from rWA-1-infected mice compared to rΔORF8-infected mice, as well as elevated RANTES/CCL5, an IFN-inducible chemokine, in ORF8-sufficient infections compared with ORF8-deficient infections (**Fig. 3A, Supplemental Fig. 2A**). Collectively, the cytokine, chemokine, and transcriptional data support activation of an IFN-I-associated inflammatory program in the ORF8-sufficient URT.

Because IFN-I signaling can intersect with epithelial secretory programs, and IFN or viral infection has been linked to mucin responses in airway epithelium (41, 42), we next tested whether IFNβ was sufficient to increase MUC5AC in mice. C57BL/6J and K18-hACE2 mice were treated daily with intranasal IFNβ, and URT samples were collected 48 hours after the initial treatment. MUC5AC protein was increased in both mouse strains following IFNβ treatment compared to PBS-treated controls (**Fig. 4A**). These findings connect IFNβ signaling to a mucus-associated effector program that may contribute to the generation of respiratory secretions.

**Figure 4.**
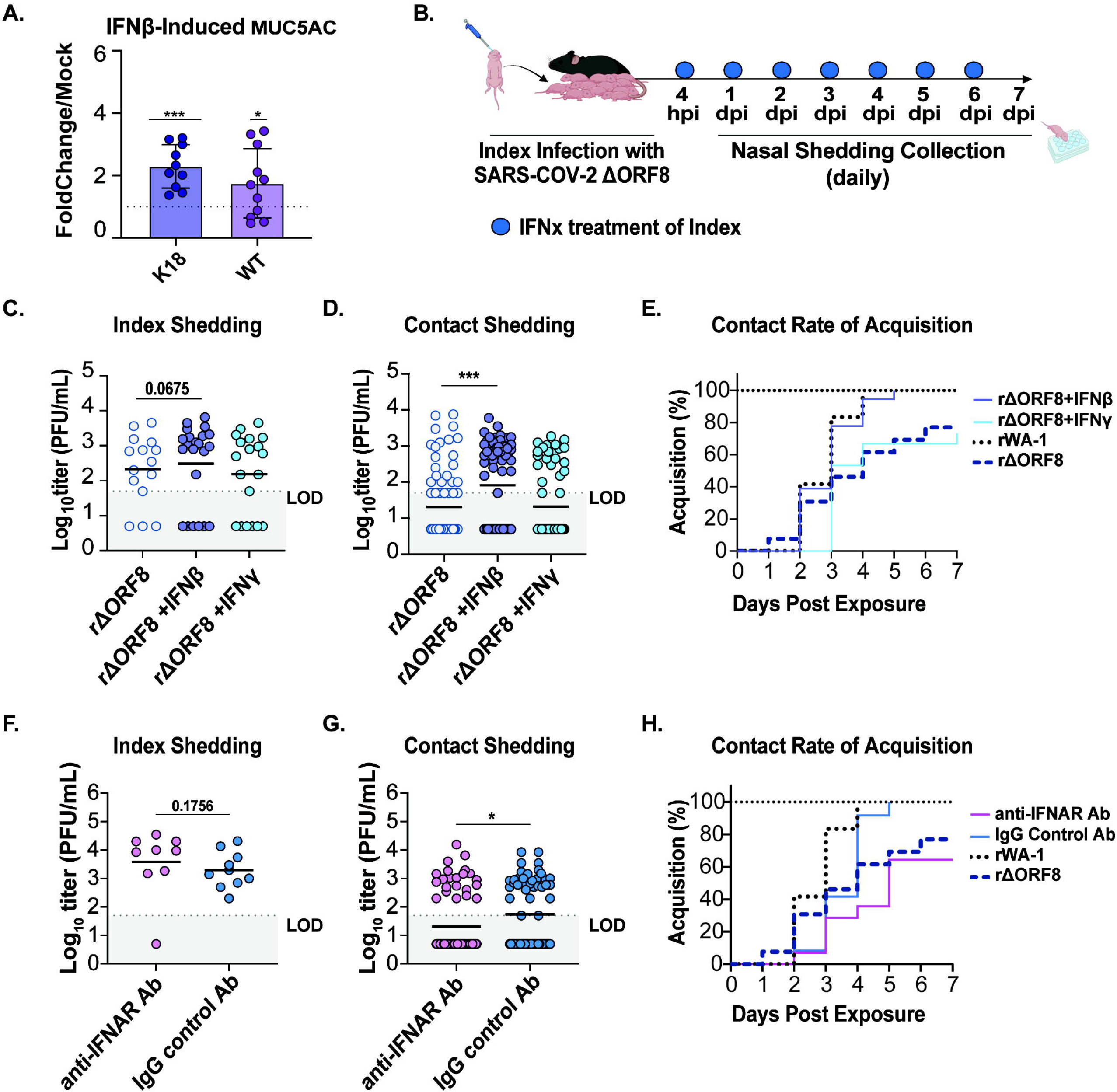
IFNβ induces MUC5AC and supports rΔORF8 shedding and transmission. (A) MUC5AC quantification in URT lavages from C57BL/6J and K18-hACE2 mice treated with IFNβ or PBS daily for 48 hours. A fold change/mock value of 1 is indicated by the dashed gray line. Each point represents the fold change/mock value for an individual mouse. (B) Schematic of IFN treatment during transmission. Index pups infected with rΔORF8 received intranasal IFNβ or IFNγ beginning 4 hpi and then daily thereafter. (C-E) Nasal shedding titers from (C) index pups and (D) contact pups collected during days 1–4. (E) Percent acquisition among exposed contacts. (F-H) IFNAR blockade experiment. Index mice were treated with α-IFNAR Ab or IgG control Ab beginning 16 hours before rWA-1 infection and then daily thereafter. Index mice were infected with rWA-1, co-housed with contacts, and nasal shedding samples were collected daily. Nasal shedding titers from (F) index pups and (G) contact pups collected during days 1–7. (H) Percent acquisition among exposed contacts.. <u>Statistics</u>. Each point in (A) (C), (D), (F), and (G) represents an individual sample. Geometric means are shown. For (A), normality was assessed using a Shapiro-Wilk test, and fold-change values were analyzed using a one-sample t test against a mock fold-change value of 1. For (C) and (D), shedding titers from all samples collected on days 1–4 were pooled by treatment group and analyzed using a Kruskal-Wallis test followed by Dunn’s multiple-comparisons test. For (F) and (G), shedding titers from all samples collected on days 1–7 were pooled by treatment group and analyzed using a Mann-Whitney U test. Values below the LOD were included and assigned a value of 5 PFU/mL, one log below the assay LOD of 50 PFU/mL. Experiments represent ≥2 independent biological replicates. Significance is denoted as *p < 0.05, **p < 0.01, ***p < 0.001, and ****p < 0.0001. Cartoons were created with BioRender.com. Graphs were generated using GraphPad Prism 10.

Collectively, the cytokine, cellular, transcriptional, and MUC5AC data identify an ORF8-associated URT inflammatory state characterized by increased cytokine and chemokine production, selective macrophage recruitment, IFN-I-associated pathway activation, and IFNβ-responsive mucin induction. These findings raised the possibility that the reduced shedding and transmission phenotype of rΔORF8 reflects loss of an IFN-I-associated URT inflammatory program rather than reduced viral RNA abundance.

### IFNβ increases URT shedding and rescues transmission during ΔORF8 infection

To test whether IFNβ was sufficient to restore the rΔORF8 shedding and transmission phenotype, rΔORF8-infected index mice were treated intranasally with IFNO or IFNγ at 4 hours post infection (hpi) and then daily thereafter **(Fig. 4B)**. IFNO supplementation restored infectious virus shedding in rΔORF8-infected index mice, whereas IFNγ did not **(Fig. 4C)**. IFNO also increased transmission to exposed contacts from 80% to 100%, while contacts exposed to IFNγ-treated index mice remained at 80% acquisition **(Fig. 4D-E)**. These findings indicate that IFNβ is sufficient to restore infectious virus shedding and rescue transmission during rΔORF8 infection.

The rescue of the rΔORF8 phenotype by IFNβ implicated IFN-I signaling as a functional contributor to host-to-host spread. To determine whether endogenous IFN-I signaling is required for efficient transmission, only index mice were treated with an antibody (Ab) blocking the type I interferon receptor (anti-IFNAR Ab) or an IgG isotype control (IgG control Ab) 16 hours before infection with rWA-1, which consistently transmits to 100% of exposed contacts. Contact mice received no antibody treatment. Anti-IFNAR Ab treatment did not significantly alter infectious virus titers measured in index nasal secretions compared with IgG Control Ab treatment **(Fig. 4F)**. However, anti-IFNAR Ab treatment reduced infectious virus recovered from exposed contacts **(Fig. 4G)** and decreased overall transmission efficiency **(Fig. 4H)**. Because IFNAR blockade was restricted to index mice, these findings indicate that endogenous IFN-I signaling in the infected host contributes to transmission, an effect captured more clearly by contact infection than by discrete measurements of index shedding. Together, IFNβ supplementation and IFNAR blockade provide complementary evidence of pathway sufficiency and physiological requirement, identifying IFN-I signaling in the infected host as a functional contributor to ORF8-dependent SARS-CoV-2 shedding and transmission.

## Discussion

Preventing viral spread remains one of the greatest challenges in pandemic control. Understanding the viral, host, and environmental factors that influence transmission is essential for predicting the emergence of new variants and designing strategies to curb viral spread. Transmission is inherently complex because it depends on multiple interacting variables, making it difficult to isolate the contribution of any single viral factor. Here, we focused on how the SARS-CoV-2 accessory protein ORF8 facilitates host-to-host spread. We found that ORF8 supports the recovery of infectious virus from the URT and nasal secretions, augments local inflammatory responses, and promotes efficient contact transmission. ORF8-deficient viruses showed reduced recovery of infectious virus from the URT without a corresponding reduction in lung titers or significant differences in genomic or subgenomic viral RNA. These findings identify the URT infectious and inflammatory environment, rather than lung infection alone, as a critical determinant of transmission. Nasal discharge, sneezing, and coughing help clear virus from the respiratory tract, but can also release pathogen-containing droplets and increase exposure of susceptible contacts (9). In this context, URT inflammation may enhance contagiousness when it promotes the expulsion of infectious virus from the infected host.

Building on this URT-centered model of contagiousness, we hypothesized that accessory proteins may enhance viral fitness by shaping the compartment from which virus is expelled rather than by directly increasing intrinsic viral replication efficiency. In our study and others, ORF8 is dispensable for SARS-CoV-2 replication in cell culture and does not produce a broad defect in lung infection (32, 33), yet its deletion reduced infectious virus recovered from URT tissues and nasal secretions (29) (**Fig.2**). This compartment-specific phenotype differs from that of low-replicating clinical isolates, including Omicron BA.1 and BQ.1.1, which shed poorly and transmitted inefficiently in the infant mouse model (29). Although viral RNA levels trended lower in rΔORF8 samples, neither genomic nor subgenomic RNA differed significantly between rWA-1 and rΔORF8 infections (**Fig.2D-E**). Although the sample size may have limited detection of modest differences in viral RNA abundance, reduction in viral RNA remained substantially smaller than the reduction in infectious virus recovered from the URT. The rΔORF8 phenotype therefore cannot be explained solely by a proportional reduction in viral RNA production.

Accordingly, we interpret ORF8 effects through matched isogenic comparisons and focus on infectious virus recovered from URT secretions as the proximal determinant of exposure dose and contact acquisition. This framework distinguishes the compartment-specific rΔORF8 phenotype from a generic reduction in viral fitness caused by unrelated attenuating mutations. The present study does not discount the importance of viral burden but identifies ORF8 as a viral determinant that selectively supports a URT inflammatory state associated with efficient shedding and spread.

The compartment-specific effect of ORF8 likely reflects its ability to modulate host immune responses in the URT. ORF8 has been shown to downregulate MHC-I (24), induce ER stress (43), and influence cytokine production (22, 26, 27). Clinical observations that naturally occurring ORF8-deletion variants are associated with milder disease and altered inflammatory responses in humans further support a role for ORF8 in shaping host inflammation (28).

Together with our findings, these observations support a model in which ORF8 enhances viral fitness by promoting a mucosal inflammatory environment associated with infectious virus shedding. Through these mechanisms, ORF8 may simultaneously limit immune recognition while promoting cytokine and epithelial secretory programs that support transmission-relevant URT inflammation. Elevated local inflammation can increase mucus production, vascular permeability, and mucociliary function in the nasal epithelium, creating conditions that may favor the release of infectious virus from the infected host. In this way, ORF8 may help sustain a URT environment that enhances viral shedding and transmissibility.

Within this ORF8-associated inflammatory environment, ORF8-sufficient viruses, including rWA-1 and rS84L+E92K, induced higher levels of inflammatory cytokines and chemokines in nasal lavages than ORF8-deficient viruses, including rΔORF8 and rQ27*. These mediators included RANTES/CCL5, MIP-1α, and MIP-1β, which are associated with leukocyte recruitment and innate immune activation. Flow cytometry of nasal lavage cells further showed increased macrophage recruitment in the URT following ORF8-sufficient infection compared to ORF8-deficient infection. Macrophages amplify local inflammation by producing cytokines and chemokines that recruit additional leukocytes and increase vascular permeability, both of which can contribute to nasal discharge. These observations identify macrophage recruitment as a potential cellular link between cytokine induction and secretion-associated URT inflammation. Macrophages may also connect this cellular inflammatory phenotype to the IFN-I pathway identified in our transcriptional and functional studies. Macrophages are a potential source of IFNβ, and ORF8-sufficient infection was associated with enrichment of IFN-I transcriptional pathways, increased IFNβ protein in nasal lavage fluid, and elevated RANTES/CCL5, an IFN-inducible chemokine (44, 45). Because IFNβ protein abundance in mucosal secretions can vary with timing, viral burden, and cellular composition, we interpret IFNβ together with downstream transcriptional and functional evidence of IFN-I pathway activation rather than as a single standalone correlate. Collectively, these findings position IFN-I signaling within a broader URT inflammatory and secretory program that links host defense to the recovery of infectious virus from nasal secretions.

The functional experiments further support this interpretation. Exogenous IFNβ supplementation increased infectious virus shedding and rescued transmission in rΔORF8-infected mice, whereas IFNγ did not. These findings indicate that IFNβ contributes to a transmission-competent URT state in the absence of ORF8. Conversely, IFNAR blockade in rWA-1-infected index mice reduced infectious virus recovered from exposed contacts and decreased overall transmission efficiency, despite not significantly changing infectious virus titers measured in index nasal secretions. Because IFNAR blockade was restricted to index mice, these data indicate that endogenous IFN-I signaling in the infected host contributes to the generation of an effective infectious exposure. This contribution was more readily detected through contact acquisition than through discrete measurements of index shedding. Together, IFNβ supplementation and IFNAR blockade distinguish pathway sufficiency from physiological requirement and identify IFN-I signaling in the infected host as a functional contributor to ORF8-dependent shedding and transmission. This finding is consistent with our previous work in the infant mouse model showing that IFN-I signaling promotes influenza A virus and Streptococcus pneumoniae shedding and transmission, supporting a broader role for IFN-I in respiratory contagiousness (46).

The differential effects of IFNβ and IFNγ should not be interpreted as evidence that the URT cannot respond to IFNγ or that IFNγ is irrelevant to respiratory virus transmission. Airway epithelial cells express functional IFNγ receptor components and respond to IFNγ signaling (47, 48). Moreover, in the infant mouse model of influenza A virus transmission, IFNγ-driven nasal inflammation promoted mucus production, viral shedding, and host-to-host transmission (49).

Under the experimental conditions used here, IFNγ did not restore rΔORF8 shedding or transmission to the levels achieved with IFNβ. It remains possible that the timing, dose, or duration of IFNγ exposure requires further optimization in this SARS-CoV-2 model to reveal a transmission-relevant phenotype. This distinction could also suggest that the inflammatory pathways that promote contagiousness are virus- and context-dependent. IFNγ may still contribute to mucosal inflammation or mucus-associated responses during SARS-CoV-2 infection, but these effects were insufficient to rescue the rΔORF8 phenotype. Other mediators, including TNF or cell death-associated inflammatory pathways, may also influence epithelial integrity, secretion, or viral release, although their contributions were not directly tested in this study. Collectively, these findings support a model in which respiratory viruses exploit distinct inflammatory circuits to enhance contagiousness, with IFNγ playing a prominent role in influenza virus transmission (49) and IFN-I representing the ORF8-linked pathway most directly supported in SARS-CoV-2. This distinction illustrates the balance respiratory viruses must achieve between immune evasion and immune exploitation to optimize transmission.

ORF8 is one of the most variable SARS-CoV-2 proteins, and this diversity may reflect changing selective pressures across viral lineages. In our model, loss or truncation of ORF8 generally impaired transmission-relevant phenotypes, whereas individual ORF8 substitutions produced distinct effects. The S84L mutation alone did not enhance transmission relative to the ancestral WA-1, which harbors serine (S) at position 84, suggesting that additional mutations, including the co-occurring D614G substitution in the Spike protein, likely contributed to the fitness of early lineages. Conversely, the Q27* mutation, which removes most of the C-terminal immunoglobulin-like domain of ORF8, reduced URT infectious titers and transmission, supporting the importance of an intact ORF8 protein for transmission-relevant function. Notably, the Alpha variant (B.1.1.7) carried the same Q27* truncation in ORF8, yet remained highly transmissible in humans, likely because mutations elsewhere in its genome compensated for the loss of ORF8 function. Similarly, the reemergence of S84L in later lineages must be interpreted within the broader genomic context of those viruses. The simultaneous evolution of multiple viral proteins makes it difficult to disentangle individual effects on transmission using epidemiologic data alone, underscoring the value of controlled *in vivo* models that test defined mutations in a consistent genetic background. Such models complement genomic surveillance and computational predictions by directly assessing how individual viral factors shape transmission-relevant phenotypes.

In summary, our findings support a model in which ORF8 promotes SARS-CoV-2 contagiousness by reshaping the URT into an inflammatory and secretory state that favors the release of infectious virus from the infected host. ORF8-dependent cytokine induction, macrophage recruitment, and IFN-I-associated signaling converge with mucus-associated responses to increase infectious virus recovered from nasal secretions and thereby enhance the exposure available for contact acquisition. These findings establish a mechanism by which a SARS-CoV-2 accessory protein can engage host mucosal responses to enhance contagiousness. Accessory protein variation, although often overlooked, can influence viral spread and should be considered alongside Spike evolution in genomic and functional surveillance. Defining how viral proteins manipulate host inflammation and shedding may inform future strategies designed not only to mitigate disease but also to limit transmission.

## Materials and Methods

### Biosafety

All work on SARS-CoV-2 infected animals was performed in the Animal Biosafety Level 3 (ABSL3) facilities at NYU Grossman School of Medicine. These facilities are registered with the local Department of Health and have passed CDC inspection. Work with SARS-CoV-2 samples derived from infected animals was performed in either ABSL3 or enhanced Biosafety Level 2 (BSL2+) facilities, as approved by the Institutional Biosafety Committee (IBC). Research in ABSL3 and BSL2+ facilities at NYU was conducted according to IBC approved standard operating procedures. Recombinant viruses were generated at the University of Maryland School of Medicine according to IBC approved protocols. Viruses were transported to NYU Grossman School of Medicine under CDC-approved import permits and material transfer agreements. Generation and transport of recombinant SARS-CoV-2 viruses were approved for Dr. Matthew Frieman by the IBC of the University of Maryland School of Medicine.

### Mice

C57BL/6J (strain # 000664, Jackson Laboratories), K18-hACE2^(+/-)^ (strain # 034860; B6.Cg-Tg(K18-ACE2)2Primn/J, Jackson Laboratories), and hACE2-KI (strain # 035000; B6.129S2(Cg)-*Ace2^tm1(ACE2)Dwnt^*/J, Jackson Laboratories) mice were maintained and bred in a conventional animal facility. Pups were housed with their dam for the duration of all experiments, and both male and female pups were used. All animal studies were conducted in accordance with the Public Health Service Policy on Humane Care and Use of Laboratory Animals, and institutional guidelines (50). All procedures were approved by the Institutional Animal Care and Use Committee (IACUC) of NYU Langone Health under protocol PROTO202200097_TR01. NYU Langone Health maintains an approved Animal Welfare Assurance with the Office of Laboratory Animal Welfare (PHS Assurance # A3317-01). All procedures were also conducted in compliance with the Biosafety in Microbiological and Biomedical Laboratories. NYU Langone Health’s animal care and use program is accredited by the Association for Assessment and Accreditation of Laboratory Animal Care International (AAALAC International).

### Cell Lines

Vero E6-TMPRSS2-T2A-ACE2 cells were obtained from BEI Resources (NR-54970) cultured in Dulbecco modified Eagle medium (DMEM, Corning) containing 4 mM L-glutamine, 4500 mg per L glucose, 1mM sodium pyruvate, and 1500 mg per L sodium bicarbonate, and supplemented with 10% fetal bovine serum (FBS, Atlanta Biologicals), 1% penicillin/streptomycin (Cytiva), 1% nonessential amino acids (Gibco), and 10 µg per mL puromycin (Gibco). Cells were maintained at 37°C with 5% CO_2_. Cells were confirmed to be mycoplasma-free.

### Viruses

Recombinant SARS-CoV-2 viruses, including rWA-1, rΔORF8, rQ27*, rS84L, rE92K, and rS84L+E92K, were generated in the Frieman laboratory at the University of Maryland as previously described (32, 33). Viral stock titers were determined by plaque assay using Vero E6-TMPRSS2-T2A-ACE2 cells.

### Virus Infection, Shedding, and Transmission

Index pups 4-7Odays of age were infected by intranasal instillation with a 1,500 PFU of SARS-CoV-2 diluted in 3μl sterile PBS and returned to the litter for the duration of the experiment.

Inoculations were performed without anesthesia to avoid direct lung deposition (35). Ratios of index to contact pups ranged from 1:6 to 1:8. Viral shedding was collected daily by dipping the nares of each mouse into viral medium (PBSO+ 0.3% bovine serum albumin [BSA]), and samples were quantified by plaque assay using Vero E6-TMPRSS2-T2A-ACE2 cells. Intra-litter transmission was assessed in naïve littermates (contacts) at 4-7Odays post-exposure (postnatal days 10-14). Animals were euthanized by CO_2_ asphyxiation in accordance with institutional guidelines. Nasal lavages were collected after euthanasia by retrograde lavage of the URT performed by flushing 300µL PBS through the trachea and collecting effluent from the nares.

Lung tissue was homogenized in PBS using a TissueLyser III (Qiagen). Homogenates were centrifuged at 8,000 rpm for 8 minutes, and cleared supernatants were collected for downstream analyses. Samples were used for plaque assays, flow cytometry, Luminex assays, RT-PCR, and RNA sequencing.

### Compound and Antibody Treatments

Interferons (IFNβ [R&D Systems, 8234-MB-010], IFNγ [R&D Systems, 169049915]) were prepared in PBS + 0.1% PBS/BSA and administered intranasally at 1ng/μL/g body weight in a 3μL volume 4 hours after infection and then once daily for 7 days. Contact mice received no treatment. IFNAR blocking antibody (InVivoMAb anti-mouse IFNAR-1 [bioXcell BE0241, clone MAR1-5A3]) or control antibody (InVivoMAb mouse IgG1 isotype control [bioXcell BE0083, clone MOPC-21] was diluted to 20 µg/pup in buffer (InVivoPure pH 7.0 Dilution Buffer [bioXcell IP0070]). Antibodies were administered intranasally in a 5µL volume 16 hours before infection and then once daily to index mice only for the duration of the experiment (7 days). Contact mice received no treatment.

### RT-PCR

Viral copy number per µL was determined via RT-qPCR using the TaqMan RNA-to-Ct One-Step RT-PCR kit (Applied Biosystems, 01204264). To quantify genomic RNA in nasal lavage samples, SARS-CoV-2 primers and probes targeting an amplicon within the nucleocapsid (N) gene of SARS-CoV-2 WA-1 were synthesized by Integrated DNA Technologies (IDT) (Forward: 5’-ATG-CTG-CAA-TCG-TGC-TAC-AA-3’; Reverse: 5’-GAC-TGC-CGC-CTC-TGC-TC-3’; Probe: 5’-/56-FAM/TCA AGG AAC/ZEN/AAC ATT GCC AA/3IABkFQ/-3’). To quantify subgenomic RNA, SARS-CoV-2 primers and probe targeting a region containing the leader transcriptional regulatory sequence (TRS) fused to the start of the envelope (E) gene coding sequence were used (Forward: 5’-CGATCTCTTGTAGATCTGTTCTC-3’; Reverse: 5’-ACAGGTACGTTAATAGTTAATAGCGT-3’; Probe: 5’-/56-FAM/ACACTAGCC/ZEN/ATCCTTACTGCGCTTCG/3IABkFQ/-3’). RNA standards for the N and E genes were generated through in vitro transcription of template plasmids using the mMESSAGE mMACHINE T7 kit (ThermoFisher Scientific, 0087523). RT-qPCR reactions were run in technical duplicate on a QuantStudio 3 Real-Time PCR System (Applied Biosystems, ThermoFisher Scientific). Experiment-specific standard curves were used for quantitation, and copy numbers were calculated using the SARS-CoV-2 reference sequence MN985325.1.

### Plaque Assay

Infectious titers were determined by plaque assay on Vero E6-TMPRSS2-T2A-ACE2 cell monolayers. Serial dilutions of virus-containing samples were prepared in DMEM (Gibco) supplemented with 1% antibiotic/antimycotic (Gibco) and incubated on cell monolayers for 1 hour at 37°C. After adsorption, a semisolid overlay (0.8% Oxoid agar [OXLP0028B, Fisher Scientific] in DMEM with 2% FBS [Atlanta Biologicals] and 1% antibiotic/antimycotic [Gibco]) was applied, and 12-well plates were incubated for 48 hours at 37°C, until plaques were visible. Cells were fixed with 10% formaldehyde overnight (Sigma, 252549), overlays were removed, plaques were stained with 0.1% crystal violet, washed with PBS, and enumerated in a blinded fashion.

### Luminex

Cytokines and chemokines in murine nasal lavage samples were measured using the Bio-Plex Pro Mouse Cytokine 23-plex, Group I Assay (Bio-Rad, 64565935). IFNα/β levels were quantified using ProcartaPlex mouse IFNα/IFNβ 2-plex (Invitrogen, 353675-008). Data were acquired on a MAGPIX^®^ System (Luminex) and quantified against assay-specific standard curves using. xPONENT^®^ software (v4.3.229.0). Values below the limit of detection (LOD) were set to half the LOD. All samples were normalized to PBS-treated controls. Clustered heatmaps were generated in R using RStudio (v2025.05.1+513).

### Mucin Immunoblot

URT lavages were collected and diluted 1:50 in Tris-buffered saline (TBS). Samples were applied to a nitrocellulose blotting membrane (Amersham Protran 0.2 µm NC, 10600094) using a Minifold II Array Blotting System (Schleicher & Schuell). Membranes were blocked overnight using 1X Carbo-Free blocking buffer (Vector Laboratories, SP-5040-125) diluted in TBS. Blots were washed with 1X TBS + 0.1% Tween-20 (Sigma, 655204). MUC5AC was detected using HRP-conjugated MUC5AC IgG1 kappa antibody (Novus Biologicals, NBP2-50390H) at 1:10,000. All solutions were sterile filtered, and blots were developed using SuperSignal West Femto Maximum Sensitivity Substrate (Thermo Scientific, 34095). Blots were imaged using a ChemiDoc Touch Imaging System (Bio-Rad), and mucin levels were quantified using ImageJ software (NIH, v1.54p).

### Flow cytometry

URT lavages were collected, centrifuged to pellet cellular fractions, and resuspended in FACS buffer (PBS, 2% FBS, 1mM EDTA). Red blood cells were lysed on ice for 20 minutes using RBC Lysis Buffer (BD Biosciences, 566349). Fc receptors were blocked for 20 minutes with anti-CD16/32, clone 93 (Biolegend, 101302). Cells were stained on ice for 20 minutes with LIVE/DEAD AF 350 (ThermoFisher Scientific, L34963) and the following antibodies: CD11b-BUV395, clone M1/70 (Biolegend, 101242), CD8a-BUV737, clone 53-6.7 (BD Biosciences, 612759), SiglecF-BV421, clone E50-2440 (BD Biosciences, 565934), CD19-Pacific Blue, clone 6D5 (Biolegend, 115526), Ly6g-BV510, clone IA8 (Biolegend, 127633), CD45.1-BV605, clone A20 (Biolegend, 110738), CD45.2-BV605, clone 104 (Biolegend, 109841), F4/80-BV650, clone BM8 (Biolegend, 123149), MHCII-BV711, clone M5/114.15.2 (Biolegend, 107643), Ly6C-BV786, clone HK1.4 (Biolegend, 128041), CD192(CCR2)-FITC, clone SA203G11 (Biolegend, 150608), CD64-PE/Cy7, clone X54-5/7.1 (Biolegend, 139313), and CD4-AlexaFluor 700, clone GK1.5 (Biolegend, 100430). Cells were fixed with 2% paraformaldehyde (Fisher Scientific, 50980494) for 10 minutes at room temperature and resuspended in FACS buffer. Samples were acquired on a ZE5 Cell Analyzer (Bio-Rad), and data were analyzed in FlowJo software (BD Biosciences, v11.2).

### Bulk RNA Sequencing and Transcriptomic Analysis

URT lavages were collected, centrifuged to pellet cellular fractions, and total RNA was extracted using the RNeasy Mini Kit (Qiagen, 74106) according to the manufacturer’s protocol, including on-column DNase digestion. Total RNA was quantified using RNA Nano Chips on an Agilent 2100 BioAnalyzer (Agilent Technologies). RNA-seq libraries were prepared using the sparQ rRNA HMR Kit (Quantabio, 95216-096) using the recommended input range of total RNA, followed by 16 cycles of PCR amplification. Final libraries were assessed using High Sensitivity DNA ScreenTape on the Agilent TapeStation 4200 (Agilent Technologies). Library concentrations were determined using Quant-iT (Thermo Fisher Scientific), and samples were pooled in equimolar ratios. The pooled libraries were sequenced as paired-end 50 bp reads on an Illumina NovaSeq X+ platform (Illumina) using a 10B 100-cycle flow cell, targeting ∼250–300 million reads per sample. FASTQ files were processed using the Seq-N-Slide RNA-Star route (10.5281/zenodo.5550459). The resulting featureCounts files and associated metadata were imported into RStudio (v2025.05.1+513) for downstream analysis using the DeSeq2 and GSEA packages.

### Statistical Analysis

Statistical analyses were performed using GraphPad Prism (v10.4.1[532]). Heatmaps and volcano plots were generated in R using RStudio (v2025.05.1+513). Each experiment included at least two independent biological replicates. For all analyses, ns = p>0.05, * = p≤0.05, ** = p≤0.01, *** = p≤0.001, **** = p≤0.0001.

For infectious virus titers, individual values are plotted as separate data points, and geometric means are shown unless otherwise indicated. Values below the limit of detection (LOD) were assigned a value of 5 PFU/mL, one log below the assay LOD of 50 PFU/mL. For shedding analyses, samples collected across the indicated sampling window were analyzed collectively by viral group rather than separately by day, unless otherwise stated in the figure legend.

Contact shedding analyses included all collected samples during the sampling window, including samples collected before acquisition, during active shedding, after shedding ceased. For comparisons between two groups, data were analyzed using a Mann-Whitney U test unless otherwise indicated. For comparisons among more than two groups, data were analyzed using a Kruskal-Wallis test followed by Dunn’s multiple-comparisons test. For Figure 4A, normality was assessed using a Shapiro-Wilk test, and MUC5AC fold-change values were analyzed using a one-sample t-test against a mock fold-change value of 1. Panel-specific statistical tests, sampling windows, reference groups, and display conventions are described in detail in the corresponding figure legends.

### Data sharing statement

All data needed to evaluate the conclusions of this study are included in the manuscript and Supplementary Materials. RNA sequencing data can be obtained from the NCBI Gene Expression Omnibus (accession # pending).

## Acknowledgements

We are grateful to members of the Ortigoza laboratory, including Mrs. Hedy L. Rocha for mice colony maintenance and breeding, and Dr. Zeineb Mhamdi for helpful discussions and technical support with viral RNA quantification. We thank Dr. Kenneth Stapleford for providing reagents for viral genome quantification and Dr. Payal Damani-Yokota for helpful discussions and technical support with flow cytometry. We also thank the NYU Genome Technology Center for performing RNA sequencing and providing technical assistance. This shared resource is partially supported by the Cancer Center Support Grant P30CA016087 at the Laura and Isaac Perlmutter Cancer Center. We thank the NYU Langone High-Containment Laboratories Core Facility for providing ABSL-3 laboratory space and technical assistance throughout the study. This project was funded in part by federal funds from the National Institute of Allergy and Infectious Diseases, National Institutes of Health, Department of Health and Human Services, under Contract No. 75N93021C00016; the NYU Grossman School of Medicine Department of Medicine; the NYU Division of Infectious Diseases and Immunology; the Vilcek Institute of Graduate Biomedical Sciences; and the Clinical and Translational Science Institute, NYU Langone Health, Grant UL1TR001445 from the National Center for Advancing Translational Sciences, National Institutes of Health.

---

**Supplemental Figure 1**

**ORF8 promotes URT infectious virus recovery in ACE2 knock-in mice.**

ACE2 knock-in (KI) pups (B6.129S2(Cg)-*Ace2^tm1(ACE2)Dwnt^*/J) were inoculated with rWA-1 or rΔORF8. Nasal shedding was collected at 2 dpi, and URT lavages and lungs were harvested immediately afterward. Infectious virus titers are shown from left to right: nasal shedding, URT lavage, and lung. <u>Statistics</u>. Each point represents an individual mouse, and geometric means are shown. Differences between groups were analyzed using a Mann-Whitney U test. Values below the LOD were assigned a value of 5 PFU/mL. Experiments represent ≥2 independent biological replicates. Significance is denoted as *p < 0.05, **p < 0.01, ***p < 0.001, and ****p < 0.0001. Graphs were generated using GraphPad Prism 10.

**Supplemental Figure 2**

**URT immune signaling and cell populations during infection.**

(A) Cytokine and chemokine data from Figure 3A visualized as a butterfly plot. (B-D) URT immune profiling at 2 dpi by flow cytometry. Panels show (B) myeloid subsets, (C) neutrophils, and (D) eosinophils for the indicated recombinant SARS-CoV-2 infections. <u>Statistics</u>. Each point represents an individual mouse. Differences between groups were analyzed using a Mann-Whitney U test. Experiments represent ≥2 independent biological replicates. Significance is denoted as *p < 0.05. Graphs were generated using RStudio and GraphPad Prism 10.

