## Supplementary figures and images for "SARS-CoV-2 ORF8 modulates the upper respiratory tract inflammatory response to facilitate transmission"

### Supplemental Figure 1

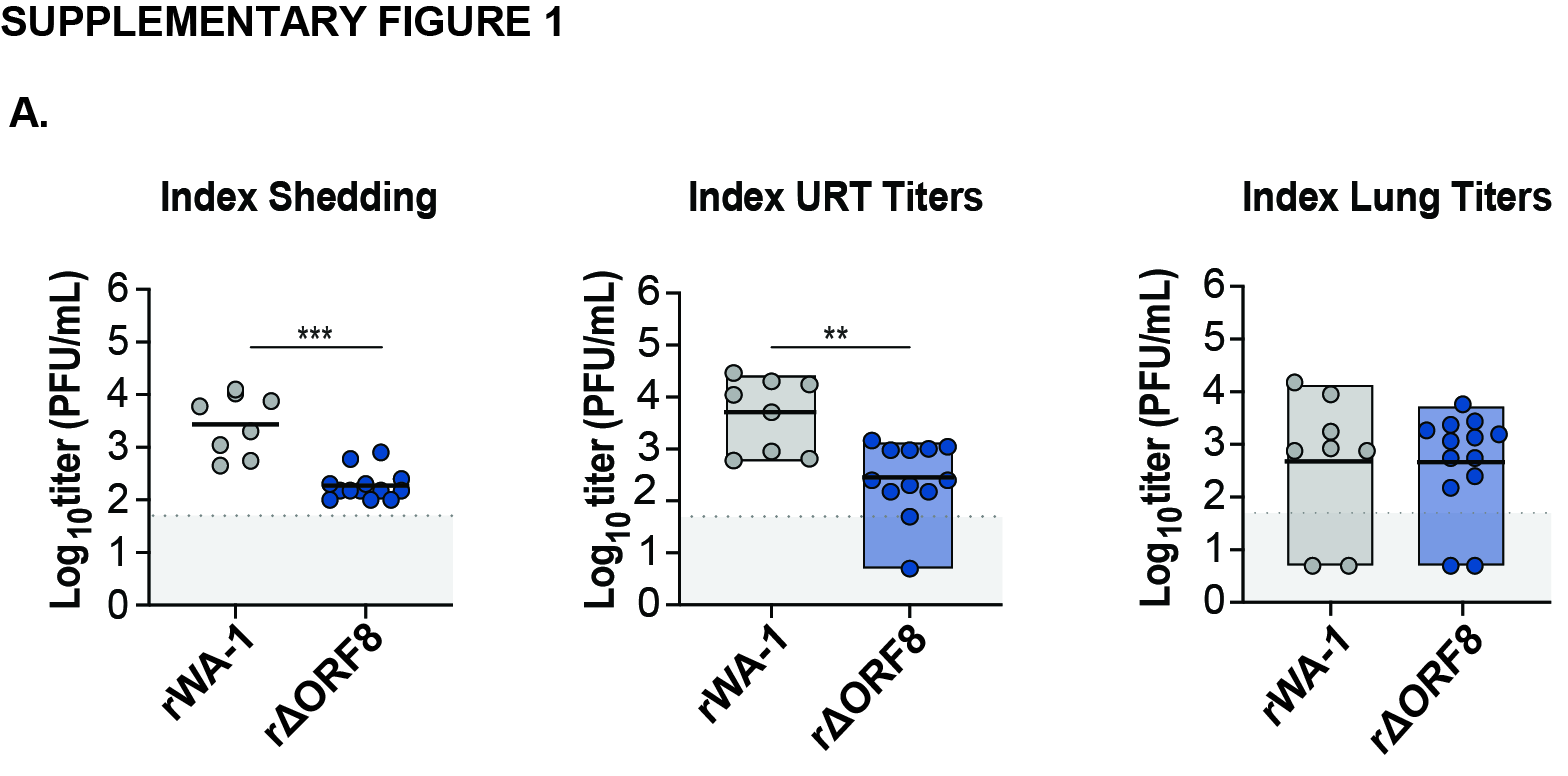

### Supplemental Figure 2

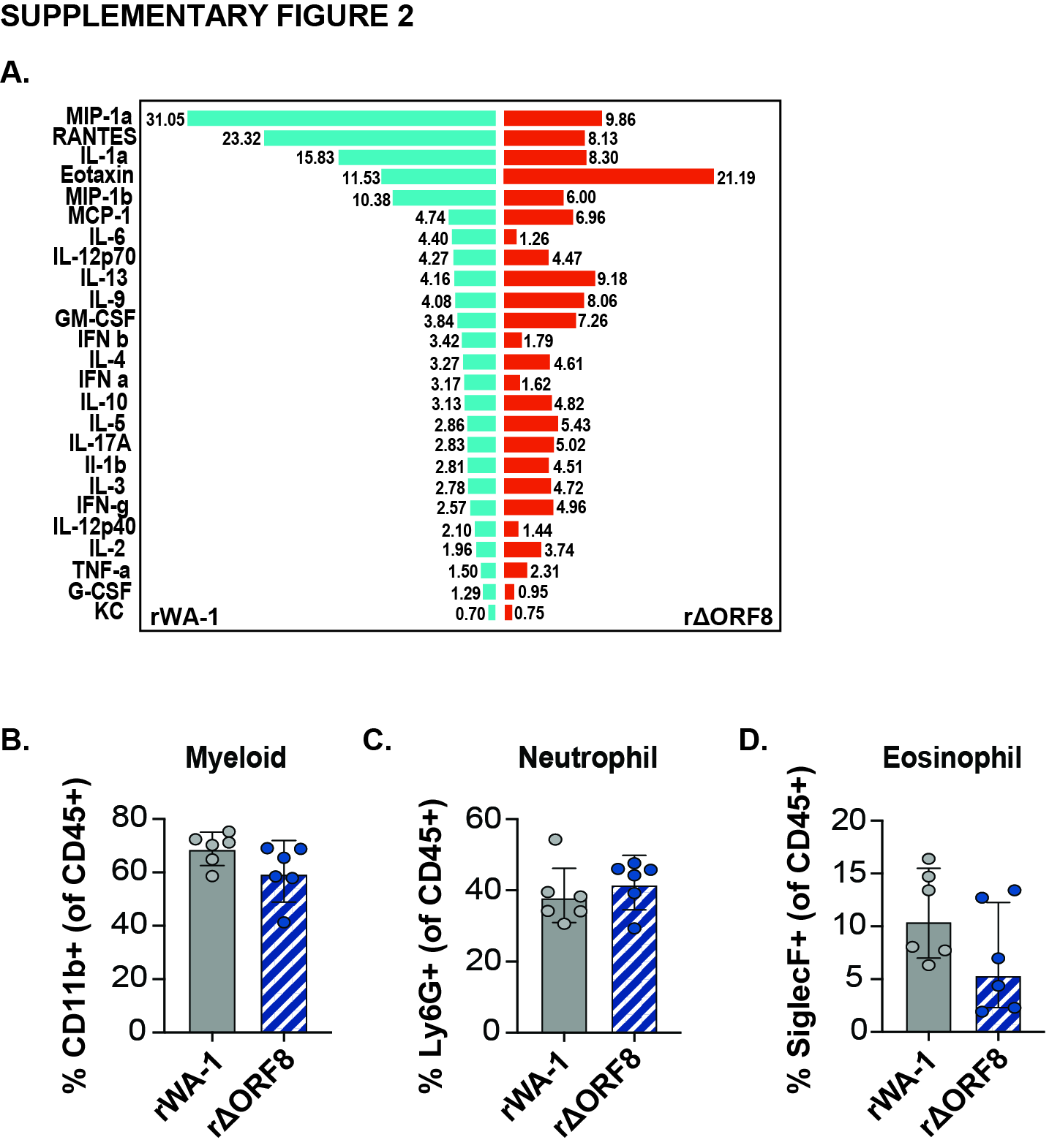
